# Pulsed taVNS-elicited pupil dilation: effects of intermixed stimulation, sham location and respiratory phase

**DOI:** 10.64898/2026.01.24.701498

**Authors:** Martin Kolnes, Sander Nieuwenhuis

## Abstract

Transcutaneous auricular vagus nerve stimulation (taVNS) may offer a powerful, noninvasive way to stimulate activity of brainstem arousal systems, including the locus coeruleus–norepinephrine (LC–NE) system. Pulsed taVNS is known to elicit pupil dilation, a marker of LC activity. However, studies reporting taVNS effects on pupil dilation have delivered taVNS and sham stimulation in separate blocks of trials or in separate sessions, hindering the integration of taVNS into rapid, event-related task designs of cognitive neuroscientists. In two experiments, we examined the effectiveness of pulsed taVNS in eliciting pupil dilation when active and sham stimulation were intermixed within the same blocks of trials, and whether these effects depend on sham location and respiratory phase. In Experiment 1 (N = 40), intermixed taVNS and (earlobe) sham pulses of 3.4 seconds produced inconclusive evidence for increased taVNS-evoked pupil dilation. In contrast, in Experiment 2 (N = 60), taVNS pulses of 1.0 seconds had a strong effect on pupil dilation, but only in a group of participants that received sham stimulation at the earlobe; this effect was abolished in a group that received sham stimulation at the upper scapha, a non-vagal control area with a similar density of sympathetic nerve fibers as the cymba concha. Furthermore, taVNS-evoked pupil dilation was not enhanced when stimulation was delivered during exhalation, as would be expected if pupil effects were mediated by the vagus nerve–nucleus tractus solitarius–LC pathway. Together, these findings show that the effect of pulsed taVNS on pupil dilation can be preserved when taVNS and (earlobe) sham are delivered in the same blocks of trials. However, the null finding with the scapha as sham location, and the absence of an enhanced taVNS effect during exhalation call into question the assumption that taVNS-induced pupil dilation is mediated by activation of the vagal afferent pathway.

## 1. Introduction

The vagus nerve serves a major role in carrying signals from various organs to the brain and back. Through its direct and indirect projections to the nucleus tractus solitarius (NTS), it modulates activity of several neuromodulatory systems (Holt, 2022), including the locus coeruleus-norepinephrine (LC-NE) system. Consequently, animal studies have found that invasive vagus nerve stimulation (iVNS) drives increases in LC firing rate and central NE release (e.g., Hulsey et al., 2017; Roosevelt et al., 2006). iVNS has been successfully used for clinical purposes in the treatment of various brain disorders (Johnson & Wilson, 2018; Wang et al., 2021). However, the method has also attracted the interest of cognitive neuroscientists who study the healthy LC-NE system and its many effects on cortical state and cognition (Poe et al., 2020).

This growing interest was further fueled by the advent of transcutaneous auricular vagus nerve stimulation (taVNS), a noninvasive, safe (Redgrave et al., 2018) and less costly alternative to iVNS that involves applying low-intensity electrical currents to the ear to stimulate the auricular branch of the vagus nerve (Farmer et al., 2021). When clinical studies confirmed that continued taVNS treatment (e.g., 30 minutes/day over a period of several months) had similar therapeutic effects as iVNS treatment (Lampros et al., 2021; Tan et al., 2023), this led to the believe that taVNS might be of major importance to cognitive neuroscientists (Van Leusden et al., 2015), by offering a powerful, noninvasive way to stimulate activity of the LC-NE system. Although this interest has not diminished, many questions remain about the stimulation parameters (e.g., intensity, duration) required to activate the LC-NE system with taVNS (Ludwig et al., 2021). In the search for optimal parameter values, researchers have mainly used the magnitude of pupil dilation as a peripheral read-out of (taVNS-induced) phasic LC activity (de Gee et al., 2017; Desbeaumes Jodoin et al., 2015; Joshi & Gold, 2020; Mridha et al., 2021). This work has established that pupil dilation, and therefore presumably phasic LC-NE activity, monotonically increases with stimulation intensity (D’Agostini et al., 2023; Ludwig et al., 2024). Moreover, a recent meta-analysis has revealed that pulsed taVNS (e.g., ∼3-second bursts) is effective at increasing pupil dilation, while more conventional protocols (continuous or 30 sec on/off stimulation) are not (Pervaz et al., 2025). These findings suggest that pulsed high-intensity taVNS (versus sham stimulation) is effective at boosting phasic LC activity.

Nevertheless, to further integrate taVNS into the rapid, event-related task designs of cognitive neuroscientists, two limitations of existing work need to be addressed. First, most previous studies reporting taVNS effects on pupil dilation have delivered taVNS and sham stimulation in separate blocks of trials (in the same session; e.g., He et al., 2022) or in separate sessions (e.g., Sharon et al., 2021). As argued by Villani et al. (2022), blocks and sessions often differ in the participant’s levels of arousal, task engagement and mental fatigue––states that have a significant impact on pupil size, and therefore confound the dependent variable of interest. Therefore, in Experiment 1 we examined whether the effects of pulsed high-intensity taVNS on pupil dilation are preserved when taVNS and sham are delivered in the same blocks of trials.

Second, most pupillometry studies that administered pulsed taVNS used stimulation durations between 3 and 5 seconds and intertrial intervals longer than 10 seconds (e.g., D’Agostini et al., 2023; Ludwig et al., 2024; Sharon et al., 2021)–– well beyond the duration of trials in typical cognitive neuroscience experiments. In Experiment 2, we considerably shortened the stimulation duration, from 3.4 to 1.0 sec.

An additional goal of Experiment 2 was to critically examine the common assumption that taVNS effects on pupil size are mediated by the pathway connecting the vagus nerve, NTS and LC. To this end, we compared two groups receiving sham stimulation at different locations (earlobe vs. scapha; Wienke et al., 2023) and examined if taVNS effects on pupil dilation were modulated by the respiratory phase at the time of stimulation (Sclocco et al., 2019).

## 2. Experiment 1

Delivery of taVNS and sham stimulation within the same block of trials, instead of in different blocks or sessions, has several advantages. First, as noted above, it prevents the effect of taVNS on pupil size from becoming confounded with pupil changes due to block-or session-related differences in psychophysiological states. For example, study participants are often more aroused and motivated and less fatigued at the start of a study or session, and these states are known to impact baseline pupil size and task-related pupil dilation (Gilzenrat et al., 2010; Hopstaken et al., 2015; Massar et al., 2018). Second, intermixed stimulation of the active and sham locations can lower desensitization due to prolonged stimulation of the same location, which may well occur in rapid task designs. And third, intermixed stimulation can possibly reduce participants’ awareness of the stimulation type (taVNS vs. sham), thereby improving the internal validity of the manipulation. A potential limitation of intermixed stimulation in combination with short intertrial intervals is that taVNS-and sham-evoked pupil dilation responses may carry over to the next trial and confound each other. Given that these pupil responses take approximately 5 seconds to return to baseline (Lloyd et al., 2023; Skora et al., 2024), mitigating these confounds requires (i) long interstimulus intervals, or (ii) statistical control of pupil dilation on the previous trial, or (iii) model-based deconvolution of the pupil signal (Wierda et al., 2012).

We are aware of only one pupillometry study that used randomly intermixed instead of block-or session-wise taVNS and sham stimulation. Villani et al. (2022) found that 3-second trains of taVNS evoked *smaller* pupil dilation compared to sham stimulation, in particular on trials with relatively small baseline pupil size. Remarkably, stimulation intensity was set just below the participants’ perceptual threshold––an approach that differs from the more established method (in clinical applications and pupillometry studies) of applying stimulation at a level that is perceptible but below the pain threshold. Furthermore, the tragus (instead of cymba concha) was chosen as active location, an option that has been criticized for lack of anatomical support (Burger & Verkuil, 2018). Thus, it remains unclear whether taVNS affects pupil dilation when it is intermixed with sham stimulation.

In Experiment 1, we used a task-free viewing condition with randomly intermixed stimulation of the active (cymba concha) and sham locations (earlobe). Other stimulation parameters closely followed the protocols used by Sharon et al. (2021) and Lloyd et al. (2023), including 3.4 seconds of stimulation per trial and matching of subjective intensities. Given that these studies found a significant taVNS effect on pupil dilation, a potential null effect in the present experiment could likely be attributed to the intermixing of stimulation types.

### 2.1. Method

#### 2.1.1. Sample

A total of 40 participants took part in the study in return for €8.50 or course credits. One participant was excluded due to excessive pupil data loss, resulting in a final sample of 39 (20.9 ± 3.5 years old, 28 female). Participants were recruited among Leiden University students through the website SONA (https://www.sona-systems.com). Exclusion criteria were neurological or cardiac disorders and use of psychoactive medication. Participants were instructed to abstain from alcohol for 24 hours and from caffeine for 3 hours prior to the start of the study. The study was approved by the Psychology Research Ethics Committee at Leiden University (S.T.-V2-4919).

#### 2.1.2. Procedure

Each participant completed one testing session lasting approximately 1 hour. Prior to the experiment, we carried out a calibration procedure to identify suitable stimulation intensities. Stimulation intensity was individually calibrated for each participant using subjective intensity ratings on a scale from 1 to 10, where 1 = no sensation, 3 = light tingling, 6 = strong tingling, and 10 = painful. Calibration began at an amplitude of 0.1 mA, increasing in steps of 0.2 mA, with a 5-seconds stimulation duration. Participants rated the sensation after each increment until they reached a rating of 9 or a maximum amplitude of 5 mA. This procedure was conducted twice—once for each stimulation location (cymba concha and earlobe). For each stimulation location, the intensity corresponding to a rating of 9 (just below painful) was used in the main experiment. Six participants reached the maximum stimulation intensity of 5 mA in the sham condition while reporting an intensity rating below 9 (mean rating = 7.2, range 6-8). Given that pupil dilation is sensitive to the perceived intensity of tactile stimuli (Ten Brink et al., 2024), including taVNS (Ludwig et al., 2024), it is important that taVNS and sham stimulation are matched in terms of subjective intensity.

Next, the eye tracker was calibrated. To minimize head movements, participants were asked to position their head on a chin rest. During the session, participants were instructed to fixate on a white fixation cross presented in the center of a grey screen.

The experiment consisted of five blocks, each containing 24 trials (12 taVNS and 12 sham), presented in a randomized order. Each trial began with a 3.4-second stimulation period, followed by a variable intertrial interval of 10–12 seconds, resultingin a total trial duration of approximately 13.4 to 15.4 seconds.

At the end of each block, participants completed a taVNS after-effects questionnaire designed to assess potential side effects (see Table S1). Note that our event-related design did not allow separate ratings for taVNS and sham.

#### 2.1.3. taVNS stimulation

To stimulate the left cymba concha and left earlobe we used two DS7A Digitimers (www.digitimer.com). To establish a good connection, the device for taVNS was equipped with a NEMOS electrode covered with cotton rings soaked in electrolyte conductive gel; the one for sham stimulation was equipped with a clip electrode covered with conductive gel. The clip electrode had a diameter of 5.5 mm and surface area of 23.8 mm^2^ per contact; the taVNS electrode was cylindrical, with a diameter of 4 mm, height of 4 mm, and effective contact area of 20-30 mm^2^ (total surface area = 50.3 mm^2^) per contact. We administered short taVNS or sham pulses, lasting 3.4 seconds per trial, characterized by a monophasic impulse frequency of 25 Hz and a pulse width of 200–300 μs. The onset and offset of each stimulation was controlled by the Psychopy experiment script.

#### 2.1.4. Pupil pre-processing steps

For pupillometry, we used a Tobii Pro eye tracker positioned 65 cm away from the participant’s eyes. Pupil size was recorded at a frequency of 60 Hz. Pupil data were pre-processed using the R package *GazeR* (Geller et al., 2020). First, the raw pupil signal was smoothed using a 100-ms moving average. Second, eye blinks were detected using a velocity-based algorithm proposed by Mathôt (2013). Eye blink onsets and offsets were identified as outliers (±2 SD) from the mean velocity profile. To avoid underestimating blink periods, these periods were extended by 100 ms in both directions. Linear interpolation was used to reconstruct the remaining data. In both experiments, qualitatively similar results were obtained after we removed any trials that contained blinks during the window from -500 ms to 2 s relative to stimulation onset.

Finally, the data were segmented by extracting samples from -0.5 s to +10 s around stimulation onset. Following Sharon et al. (2021), pupil size was converted to percentage change values relative to a 500-ms baseline before the stimulation onset using the formula: ((x - baseline)/ baseline) * 100). We also confirmed that the results remained the same when applying subtractive baseline correction, which is usually the preferred method (Mathôt et al., 2018). Two criteria were used for exclusion of data. First, we checked whether any participants had more than 50% missing raw data. One participant was excluded based on this criterion. Second, trials with more than 50% missing raw data were excluded. This led to an average loss of 1.9% of the 120 taVNS trials and 1.8% of the 120 sham stimulation trials.

#### 2.1.5. Statistical Analysis

Before the start of the study, we decided to stop data collection when the evidence for (BF_10_) or against (BF_01_) the main hypothesis (i.e., larger pupil dilation to taVNS than to sham) would reach a Bayes factor of 8 (i.e., “strong evidence”), with a maximum of 40 participants due to resource limitations (Lakens, 2022).

The analyses closely followed Sharon et al. (2021) and Lloyd et al. (2023). To analyze the pupil time series, we first applied non-parametric Wilcoxon signed-rank tests across all time points within the first 10 seconds of the trial, starting from the onset of stimulation, with data downsampled to 30 Hz. To address the issue of multiple comparisons, alpha levels (set at 0.05) were adjusted using false discovery rate correction (FDR; Benjamini & Hochberg, 1995). To quantify the effects of taVNS on pupil dilation summary measure, we conducted frequentist and Bayesian paired-samples *t*-tests using R (version 4.0.3) with the *BayesFactor* package, employing the default Cauchy prior (*r* = 0.707) in JASP (version 0.9). We also quantified a summary measure: the mean pupil size in a 1-s window centered on the pupil dilation peak, as determined from the grand-average pupil waveform in the taVNS condition. This window was then applied to all individual participants. All reported correlations are Spearman’s rank correlation coefficients.

### 2.2. Results

#### 2.2.1. Stimulation intensity

First, we tested whether there was a difference in current intensities between taVNS and sham stimulation. In line with previous studies (Lloyd et al., 2023; Sharon et al., 2021), individual stimulation intensities were higher for sham stimulation (*M*_*sham*_ = 3.1 mA; range: 1.3 to 5.0 mA) compared to taVNS (*M*_*taVNS*_ = 1.7 mA; range: 0.6 to 4.1 mA; *p* < .001).

#### 2.2.2. taVNS effects on pupil size

Figure 1A shows the pupil waveforms elicited by taVNS and sham stimulation. Pupil size reached a peak of 2.85 ± 0.19% above baseline at 1500 ms after taVNS onset, while sham stimulation reached a peak of 1.99 ± 0.19% at 1516 ms. A sample-by-sample analysis yielded no significant differences between the stimulation types (*p* > .05, two-sided Wilcoxon signed-rank test, FDR-corrected across all timepoints).

**Figure 1.**
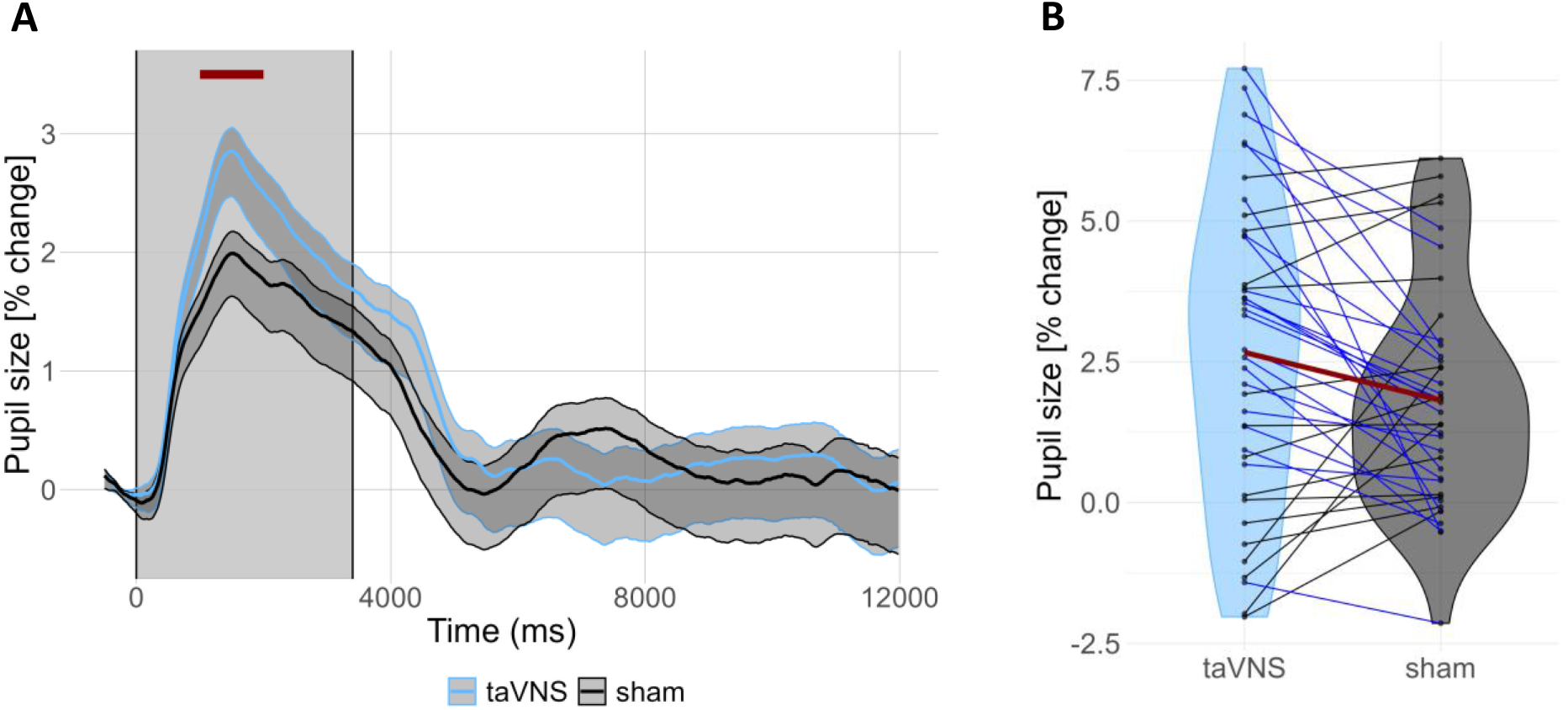
A) Grand-average pupil waveforms (percent change) relative to the 500-ms baseline period preceding stimulation onset. Shaded areas represent ±1 SEM (blue for taVNS, grey for sham). The grey-shaded rectangle denotes the stimulation period (3.4 s). The horizontal red line highlights the interval around the taVNS-evoked pupil peak (1500 ms ± 500 ms) that we used to compute pupil dilation magnitude. B) Vinyl plot showing the magnitude of pupil dilation in response to taVNS and sham stimulation. Blue lines indicate participants with the expected effect (taVNS > sham, n = 24), while black lines indicate those with the opposite effect (sham > taVNS, n = 15). The red line represents the average for both stimulation types. Like Sharon et al. (2021) and Lloyd et al. (2023), we found large between-subject variability in the magnitude of pupil dilation responses, with some participants showing little or no response, or pupil constriction.

The magnitude of pupil dilation, defined as the mean pupil size in a 1-second window centered on the taVNS-evoked pupil peak, was significantly larger for taVNS than sham (*p* = .029; Figure 1B). However, a Bayesian *t*-test revealed only “anecdotal evidence” (Lee & Wagenmakers, 2014) in favour of the alternative hypothesis, *BF*_*10*_ = 1.46.

To assess potential relationships between objective stimulation intensity and pupil dilation magnitude, we correlated these measures across participants, separately for each stimulation type. We found no significant correlation for taVNS (*ρ* =.09, *p* = .58) and sham (*ρ* = .23, *p* = .16).

### 2.3. Discussion

In Experiment 1, our design was closely based on two studies that previously reported significant effects of pulsed taVNS (versus sham) on pupil dilation (Lloyd et al., 2023; Sharon et al., 2021), except we intermixed taVNS and sham trials instead of presenting them in separate blocks. Despite our larger sample size (N = 39) than the sample size in these previous studies (Lloyd: N = 29; Sharon: N = 24), we found inconclusive evidence for an effect of taVNS on pupil dilation. While the frequentist test yielded a significant effect (*p* = .028), the Bayesian *t*-test revealed only weak support for the alternative hypothesis (BF_10_ = 1.46). Although these results suggest that intermixing taVNS and sham trials leads to less robust effects of taVNS on pupil dilation, possibly due to carry-over effects, we decided to perform a follow-up experiment in which we shortened the stimulation duration.

## 3. Experiment 2

In Experiment 2, we introduced three changes to the study design, while preserving the intermixed presentation of taVNS and sham trials: a shorter stimulation duration, an additional sham location, and the inclusion of a respiration measure.

First, we made a minor adjustment to the study design to examine whether shorter bursts of stimulation might have a more robust effect on pupil size. As noted above, a recent meta-analysis found that pulsed taVNS is effective in enhancing pupil dilation (Pervaz et al., 2025). However, there is still considerable variability in the stimulation durations used across those studies, ranging from 0.5 (Wienke et al., 2023) to 5 seconds (D’Agostini et al., 2023). In Experiment 2, we opted for a 1-second stimulation period instead of 3.4 seconds. A previous study using this duration in combination with blockwise presentation of active and sham trials reported a highly robust taVNS effect on pupil dilation (Skora et al., 2024). The reduced stimulation duration also constituted another step toward the final integration of taVNS into rapid, event-related cognitive task designs, including synchronization of taVNS and stimulus presentation.

A second change in Experiment 2 was that we compared two groups receiving sham stimulation at different locations, contrasting the earlobe with an alternative sham location (the scapha). While the vast majority of taVNS studies has used the earlobe as sham stimulation location (Farmer et al., 2021), Cakmak (2019) has criticized this choice, pointing out that the lower parts of the outer ear, including the earlobe, contain fewer sympathetic nerve fibers than the cymba concha. This is problematic because activation of sympathetic nerves can modulate sensory signals and hence result in pupil size changes through pathways other than the vagus nerve-NTS-LC pathway (Cakmak, 2019; Nieuwenhuis et al., 2011). To address this concern, a few research groups (Keute et al., 2018; Liu et al., 2016; Wang et al., 2022; Wienke et al., 2023) have chosen to apply sham stimulation to the superior scapha. The scapha, the elongated groove between the outer rim and the inner ridge of the external ear auricle, has a similar density of sympathetic nerve fibers as the cymba concha, but is not innervated by the auricular branch of the vagus nerve. Therefore, we compared groups of participants that received sham stimulation of the earlobe or the superior scapha. If taVNS would elicit larger pupil dilation than stimulation of the earlobe, but not the scapha, this would cast doubt on the involvement of the vagus nerve-NTS-LC pathway.

Third, we monitored participants’ respiration throughout the experiment. Sclocco et al. (2019), using ultra-high-field fMRI, found that effects of taVNS on BOLD activation in the NTS, LC and other brainstem nuclei was larger when taVNS was delivered during exhalation compared to inhalation. A possible explanation is that neuronal firing in the NTS, which mediates effects of the vagus nerve on LC activity, is suppressed during inhalation (Miyazaki et al., 1998). Accordingly, if we would find that taVNS elicits a larger pupil response when taVNS is delivered during exhalation, that would provide evidence for the involvement of the vagus nerve-NTS-LC pathway. As a secondary analysis, we examined if pupil size varied across the respiratory cycle, irrespective of stimulation (Schaefer et al., 2025).

### 3.1. Method

The methods of Experiment 2 were identical to those of Experiment 1, with the following exceptions.

#### 3.1.1. Sample

A total of 60 participants took part in the study. None of them had participated in Experiment 1. The experiment used a between-subjects design based on the sham location. One participant was excluded due to excessive pupil data loss. In one group, the earlobe was used as the sham location (n = 30; age: 20.7 ± 3.5 years; 28 female). In the other group, the scapha was used as the sham location (n = 29; age: 20.6 ± 2.6 years; 24 female).

#### 3.1.2. Procedure

At the start of the experiment, before the calibration procedure, a respiration belt was placed around the participant’s chest or abdomen, worn over close-fitting clothing. For one group, the sham electrode was placed on the left earlobe; for the other group, it was placed on the upper scapha of the left ear (Wienke et al., 2023). We conducted the same calibration procedure to identify suitable stimulation intensities. Three participants in the earlobe group and three in the scapha group reached 5 mA in the sham condition while reporting intensity ratings below 9 (mean_earlobe_ = 6.7, range_earlobe_ 5-8; mean_scapha_ 5.7, range_scapha_ 5-6). Only one participant in the taVNS condition (earlobe group) reported a lower rating (7).

The experiment consisted of 140 trials divided into five blocks, each containing 28 trials (14 taVNS and 14 sham), presented in a randomized order. Each trial began with a 1-second stimulation period, followed by a variable intertrial interval of 10–12 seconds, resulting in a total trial duration of approximately 11 to 13 seconds. At the end of each block, participants completed the taVNS after-effects questionnaire designed to assess potential side effects (see Table S2).

#### 3.1.3. Pupil pre-processing

The preprocessing of the pupil data led to an average loss of 1.0% of the 140 taVNS trials and 1.0% of the 140 sham stimulation trials. As in Experiment 1, pupil size was converted to percentage change values relative to a 500-ms baseline preceding stimulation onset. We also confirmed that the results remained unchanged when applying subtractive baseline correction. One participant was excluded from the final sample due to insufficient pupil data.

##### 3.1.4. Respiration pre-processing steps

The raw respiration signal was recorded using a BIOPAC MP-150 system (BIOPAC Systems, Inc., Goleta, CA) at a sampling rate of 50 Hz. Pre-processing was performed using the NeuroKit2 Python package (Makowski et al., 2021), which incorporates the following steps. First, the raw signal was preprocessed using a second-order Butterworth bandpass filter with a cutoff frequency range of 0.05–3 Hz to remove high-frequency noise and low-frequency trends. Second, peak detection was performed based on the method described by Khodadad et al. (2018), which optimizes the detection of respiratory cycles. Third, the respiratory volume per time was computed using the approach outlined by Harrison et al. (2021), which employs a Hilbert-based method for processing respiratory time series. Finally, respiratory cycles were delineated to extract key features, including respiratory rate, amplitude, and respiratory phase (inhalation and exhalation), identified from peaks and troughs in the signal. One participant from the scapha group was excluded from respiration analysis due to insufficient data.

#### 3.1.5 Statistical Analysis

Before the start of the study, we decided to stop data collection when the evidence for (BF10) or against (BF01) the main hypothesis (i.e., larger pupil dilation to taVNS than to sham) would reach a Bayes factor of 8 (i.e., “strong evidence”), with a maximum of 30 participants per group due to resource limitations (Lakens, 2022). Data collection was limited to a maximum of 30 participants per group due to resource limitations.

All analyses were done in R using the *afex* and *lme4* packages. To examine differences in current intensities between the taVNS and sham stimulation, we performed a 2 (group: earlobe, scapha) × 2 (stimulation type: taVNS, sham) mixed ANOVA. To assess stimulation effects on pupil dilation, stimulation-evoked pupil responses were quantified as the mean pupil dilation around the taVNS-evoked peak (1433 ms ± 500 ms). For each trial, we extracted the respiratory phase (bin number) at stimulation onset and the corresponding pupil dilation magnitude. These data were analyzed using an 18 (respiratory bin) × 2 (group) × 2 (stimulation type) mixed ANOVA.

The respiration analyses were divided into two sets. First, we sought to replicate the approach used by Schaefer et al. (2025), who examined how pupil size changes throughout the respiratory cycle. Second, we examined whether respiratory phase modulated the effect of taVNS and sham stimulation on the pupil dilation response. To analyse these effects of the respiratory cycle, each trial was divided into 18 evenly spaced respiratory bins, each covering 20 degrees of the respiratory cycle. Because the inhalation phase is typically shorter than the exhalation phase, the inhalation bins contained a smaller amount of data.

The first set of respiration analyses focused on examining how pupil size varied over the 18 bins of the respiratory cycle. For this analysis, we used continuous pupil data without baseline correction, following the approach of Schaefer et al. (2025). To assess whether mean normalized (z-scored) pupil size differed across the respiratory cycle—and whether there were interactions with stimulation type (taVNS vs. sham) or group (earlobe vs. scapha)—we conducted an 18 × 2 × 2 mixed ANOVA. Furthermore, like Schaefer et al. (2025), we performed non-parametric permutation tests to determine for each respiratory bin whether the mean normalized pupil size significantly deviated from zero. First, we calculated a one-sample *t*-statistic for each of the 18 respiratory bins. Second, to generate a null distribution of *t*-values, we conducted 10,000 permutations per bin, randomly flipping the signs of the observations in each permutation. This procedure preserved the magnitude and variability of the original data while simulating datasets consistent with the null hypothesis that the mean normalized pupil size is zero. Finally, the observed *t*-statistic for each respiratory bin was compared against the corresponding null distribution of maximum absolute *t*-statistics (Schaefer et al., 2025). Family-wise error-corrected *p*-values were calculated as the proportion of permuted maximum *t*-statistics that were greater than or equal to the observed absolute *t*-statistic.

In a second set of analyses, we examined whether respiratory phase modulated taVNS-and sham-evoked changes in pupil size by assessing the main effect and interaction effects of respiratory bin in our 18 × 2 × 2 mixed ANOVA. As a follow-up analysis, we conducted non-parametric permutation tests to assess, for each respiratory bin, whether pupil responses were systematically larger or smaller than the participant’s average evoked response. To ensure comparability with the first set of respiration analyses, pupil data were z-scored within participant and baseline-corrected prior to analysis. Next, pupil dilation magnitudes were mean-centered within participant prior to testing, such that values reflected deviations from the individual’s overall response. Null distributions of the maximum absolute *t*-values were generated while collapsing across groups and stimulation types.

To illustrate the effects of the respiratory cycle, we created polar plots and polar histograms in R. First, we calculated the average normalized pupil size for each participant and each of the 18 discrete bins. These binned averages were then used to calculate a weighted circular mean direction for each participant, where the weights correspond to the average pupil size per bin and the angles correspond to the center of each respiratory bin in radians. The weighted circular mean was computed using the standard formula for circular data:

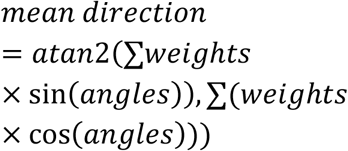

Finally, we visualized the data by plotting participant-level traces, group-level averages, and a polar histogram of the distribution of participants’ circular mean directions, providing a summary of where pupil size tended to peak across the respiratory cycle.

### 3.2. Results

#### 3.2.1. Stimulation intensity

The stimulation intensities in the earlobe group were *M*_*sham*_ = 3.4 mA (range: 2.1 to 5.0 mA) and *M*_*taVNS*_ = 1.8 mA (range: 0.6 to 5.0 mA). Those in the scapha group were *M*_*sham*_ = 3.0 mA (range: 0.7 to 5.0 mA) and *M*_*taVNS*_ = 2.0 (range: 0.6 to 4.5 mA). As in Experiment 1, stimulation intensities were higher for sham stimulation, *F*(1, 57) = 78.65, *p* < .001, *generalized η^2^* = .284, but the difference between sham and taVNS was less pronounced in the scapha group, as indicated by a significant interaction effect, *F*(1, 57) = 4.89, *p* = .031, *generalized η^2^* = .024.

#### 3.2.2. taVNS effects on pupil size

Figure 2A shows the pupil waveforms elicited by taVNS and sham stimulation in Experiment 2, separately for the earlobe and scapha groups. In the earlobe group, pupil size reached a peak of 5.68 ± 0.20% above baseline at 1433 ms after taVNS onset, while sham stimulation reached a peak of 3.54 ± 0.19% at 1533 ms. In the scapha group, pupil size reached a peak of 4.77 ± 0.22 % above baseline at 1433 ms after taVNS onset, while sham stimulation reached a peak of 4.27 ± 0.21% at 1433 ms. Although the pupil response elicited by taVNS was numerically larger in the earlobe group than in the scapha group, this difference was not significant (Δ = 0.91%, *p* = .40).

**Figure 2.**
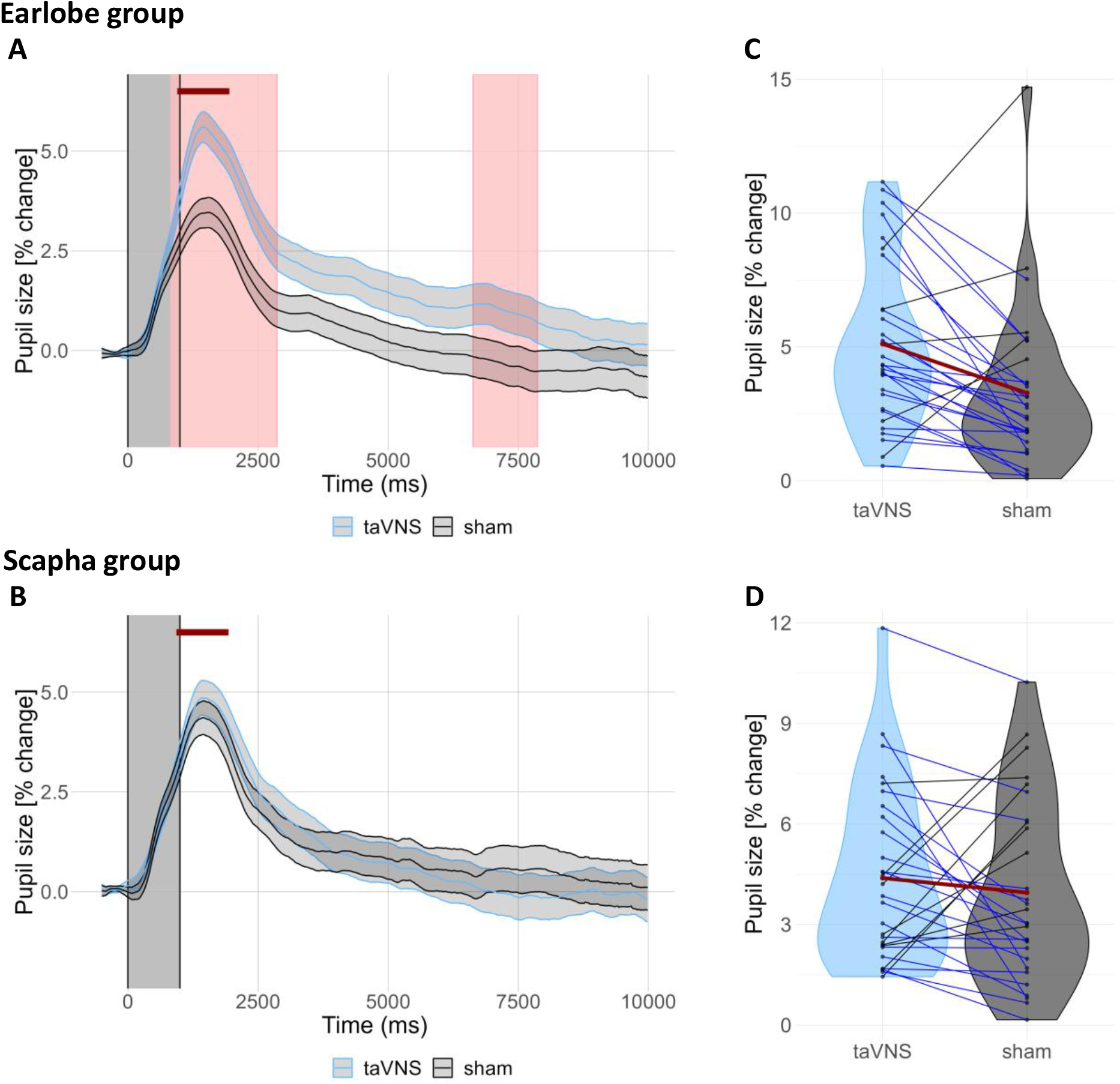
A) Grand-average pupil waveforms (percent change) relative to the 500-ms baseline period preceding stimulation onset for the earlobe group. Shaded areas represent ±1 SEM (blue for taVNS, grey for sham). The grey-shaded rectangle denotes the stimulation period (0-1 s). The horizontal red line highlights the interval around the taVNS-evoked pupil peak (1433 ms ± 500 ms) that we used to compute pupil dilation magnitude. B) Same as A, but for the scapha group. C) Violin plot depicting the magnitude of pupil dilation in response to taVNS and sham stimulation. Blue lines indicate participants with the expected effect (taVNS > sham, n = 25), while black lines indicate those with the opposite effect (sham > taVNS, n = 5). The red line represents the overall average for both stimulation types. D) Same as C, but for the scapha group. Here, 20 participants show the expected effect whereas 9 participants show the opposite effect.

Sample-by-sample analyses in the earlobe group revealed significant differences between the stimulation types from 833 to 2867 ms and from 6617 to 7883 ms after stimulation onset (*p*s < .05, two-sided Wilcoxon signed-rank test, FDR-corrected across all time points). In contrast, sample-by-sample analyses in the scapha group revealed no significant differences between the stimulations (all *p*s > .05).

As in Experiment 1, we also compared the mean pupil dilation magnitudes in 1-second windows centered on the pupil peaks. A mixed ANOVA (Table 1) revealed a significant main effect of stimulation type, *F*(17, 7872.2) = 42.37, *p* < .001, *η*^*2*^_*p*_ = .013, indicating greater pupil dilation with taVNS compared to sham stimulation (Figure 2C), and a significant interaction between group and stimulation type, *F*(17, 7872.2) = 7.88, *p* = .005, *η*^*2*^_*p*_ = .001. Pairwise comparisons collapsing across respiratory bins revealed strong Bayesian evidence for a larger pupil dilation response to taVNS than to sham in the earlobe group, *t*(29) = 3.31, *p* = .002, *BF*_*10*_ = 14.86, but moderate evidence for the null hypothesis in the scapha group, *t*(28) = 0.80, *p* = .43, *BF*_*01*_ = 3.77.

**Table 1.** ANOVA examining the effects of respiratory phase bin, group and stimulation type on stimulation-evoked pupil dilation magnitude.

| Effect | *df* | *F* | $\eta^2_p$ | *p* |
| --- | --- | --- | --- | --- |
| Respiratory bin (1 to 18) | 17, 7881.5 | 6.09 | 0.013 | < .001 |
| Group (earlobe vs. scapha) | 1, 56.2 | 0.004 | $7 \times 10^{-5}$ | .95 |
| Stimulation type (taVNS vs. sham) | 1, 7872.2 | 42.37 | 0.005 | < .001 |
| Respiratory bin × Group | 17, 7881.5 | 1.01 | 0.002 | .44 |
| Respiratory bin × Stimulation type | 17, 7881.1 | 1.35 | 0.003 | .15 |
| Group × Stimulation type | 1, 7872.2 | 7.88 | 0.001 | .005 |
| Respiratory bins × Group × Stimulation type | 17, 7881.1 | 0.71 | 0.002 | .90 |

Note. Analyses were conducted on trial-level data using a linear mixed-effects model to account for the hierarchical structure of trials nested within participants. Degrees of freedom were estimated using the Satterthwaite approximation.

To assess potential relationships between objective stimulation intensity and pupil dilation magnitude, we correlated these measures across participants, separately for each group and stimulation type. For the earlobe group, we found no significant correlation for taVNS (*ρ* = .06, *p* = .77) and for sham (*ρ* = -.11, *p* = .58). For the scapha group, there were also no significant correlations for taVNS (*ρ* = – .20, *p* = .29) and for sham (*ρ* = –.23, *p* = .22).

#### 3.2.3. Effects of respiratory phase on pupil size

Figure 3 shows the pupil size changes over the course of the respiratory cycle. In line with Schaefer et al. (2025), we found a significant main effect of respiratory phase on pupil size, *F*(17, 2016) = 19.11, *p* < .001, indicating that pupil size varied across the respiratory cycle (Figure 3A). However, no significant interactions were observed between respiratory phase, group and stimulation type (Table 2).

**Table 2.** ANOVA examining the effects of respiratory phase type, group and stimulation type on pupil size.

| Effect | *df* | *F* | $\eta_p^2$ | *p*-value |
| --- | --- | --- | --- | --- |
| Respiratory bin (1 to 18) | 17, 2016 | 19.11 | 0.139 | < .001 |
| Group (earlobe vs scapha) | 1, 2016 | 0.04 | $2 \times 10^{-5}$ | .85 |
| Stimulation type (taVNS vs sham) | 1, 2016 | 33.67 | 0.016 | < .001 |
| Respiratory bin × Group | 17, 2016 | 0.43 | 0.004 | .98 |
| Respiratory bin × Stimulation type | 17, 2016 | 0.16 | 0.001 | .99 |
| Group × Stimulation type | 1, 2016 | 54.51 | 0.026 | < .001 |
| Respiratory bin × Group × Stimulation type | 17, 2016 | 0.40 | 0.003 | .99 |

Note. Analyses were conducted on aggregated data using a linear mixed-effects model to account for the hierarchical structure of conditions nested within participants. Degrees of freedom were estimated using the Satterthwaite approximation.

**Figure 3.**
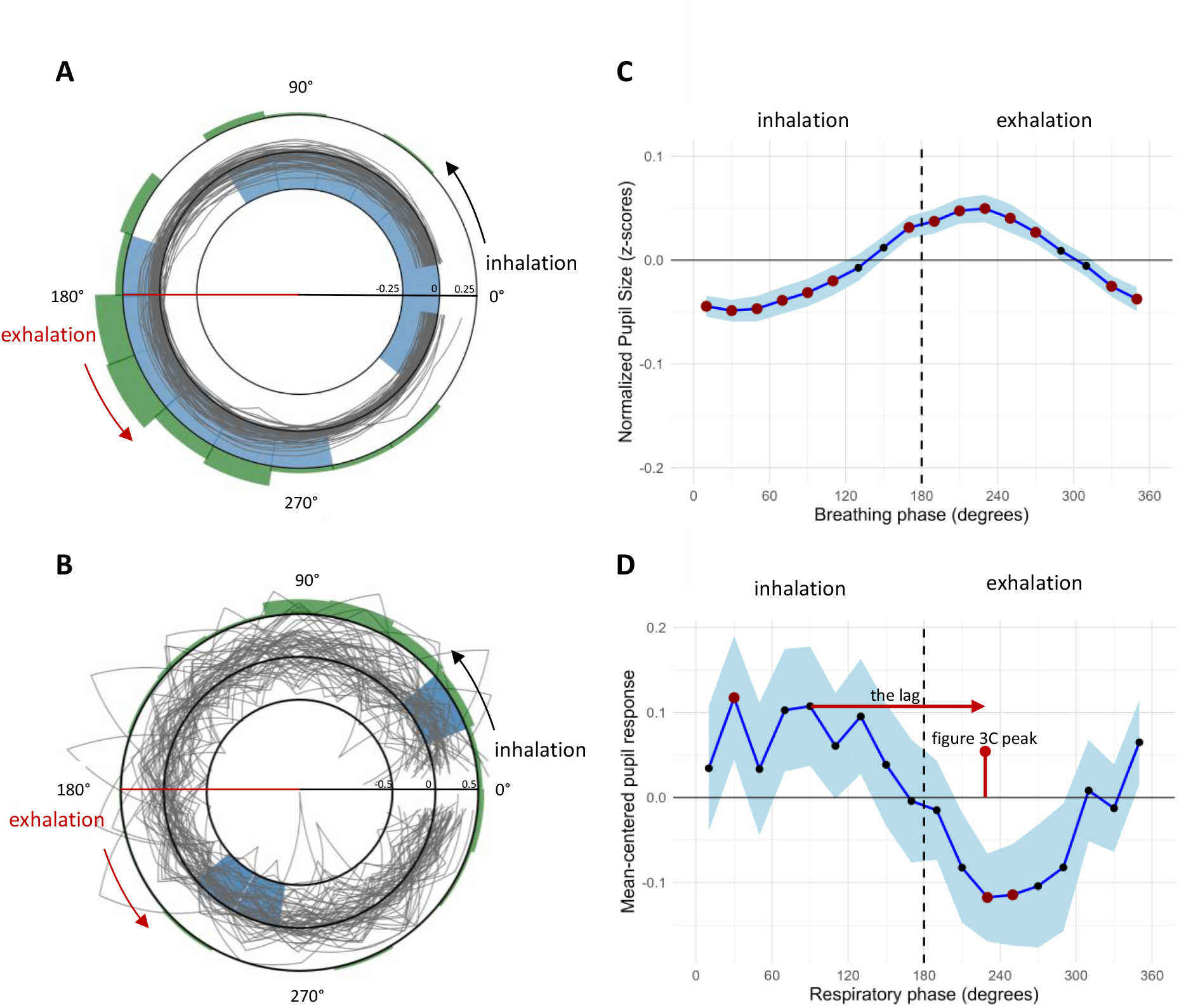
(A) Polar plots illustrating participants’ pupil size throughout the respiratory cycle. Inhalation covers 0° to 180°, and exhalation covers 180° to 360°. Each gray line represents the normalized (z-scored) pupil size pattern of an individual participant. The middle circle represents the pupil size z-score of 0. The inner and outer black circles correspond to pupil size z-scores of -0.25 and 0.25, respectively. Light blue shading marks bins where pupil size significantly differed from zero according to permutation testing. Shaded bins between the middle and outer circles indicate increased pupil size, whereas shaded bins between the middle and inner circles indicate decreased pupil size. The green polar histogram on the outside of the polar plot depicts the proportion of participants’ circular mean directions pointing towards each respiratory phase bin (the histogram sums to 1). (B) Same as (A) but for the magnitude of participants’ (mean-centered) pupil dilation in response to stimulation as a function of the respiratory phase at stimulation onset. The inner and outer black circles in the polar plot correspond to pupil size scores of -0.5 and 0.5, respectively. (C) Line plots showing the mean normalized pupil size for each respiratory phase bin. Blue shading reflects 95% confidence intervals. Red dots indicate bins in which pupil size was significantly smaller or larger than zero. (D) Same as (C) but for the magnitude of participants’ (mean-centered) pupil dilation in response to stimulation as a function of the respiratory phase at stimulation onset. The horizontal red line indicates the temporal lag relative to the peak observed in panel C.

To statistically assess the direction of the effect, we examined for which respiratory phase bins mean normalized pupil size was smaller or larger than zero. Permutation testing revealed that pupil size was significantly smaller during the end of the exhalation phase (320°-360°) and during most of the inhalation phase (0°-120°; Figure 3B). Conversely, pupil size was significantly larger during the end of the inhalation phase (160°-180°) and during the first half of the exhalation phase (180°-280°). This pattern is similar to that reported by Schaefer et al. (2025), who suggested it reflects the activity of the pre-Bötzinger complex, a medullary microcircuit that generates the breathing rhythm and projects to NE neurons in the LC that regulate pupil-linked arousal (Del Negro et al., 2018).

#### 3.2.4. Effects of respiratory phase on stimulation-evoked pupil dilation

### 3.3. Discussion

In Experiment 2, we used a shorter stimulation duration than in Experiment 1, compared two groups receiving sham stimulation at different locations (earlobe vs. scapha), and measured respiration to examine the relationship between respiratory phase and our pupil measures. We summarize the key findings. First, when taVNS was intermixed with sham stimulation of the earlobe, we found strong Bayesian evidence for an effect of taVNS on pupil dilation. Indeed, the evidence was considerably stronger than in Experiment 1, suggesting that taVNS was more effective with the 1-second stimulation duration used in Experiment 2 (earlobe group: Δ_pupil_ = 2.14%, *BF*_*10*_ = 14.86) than with the 3.4-second stimulation duration used in Experiment 1 (Δ_pupil_ = 0.86%; *BF*_*10*_ = 1.46), although the experiment × stimulation type interaction in a between-experiment ANOVA was not statistically significant (*p* = .14).

Second, we found moderate evidence against an effect of taVNS when it was compared with sham stimulation of the scapha. This seems inconsistent with Wienke et al. (2023), who reported significantly larger pupil dilation after pulsed stimulation of the cymba concha than after stimulation of the scapha. However, a limitation of this study is that the stimulation intensity was set to 2 mA for both taVNS and sham. In this case, matching should have taken into account differences in nerve density and sensitivity between the cymba concha and the scapha. Our stimulation intensitiy levels in the scapha group (M_taVNS_ = 1.9 mA, M_sham_= 3.0 mA) suggest that the scapha is less sensitive to stimulation than the cymba concha. This could explain why in Wienke’s study, the same stimulation intensity led to smaller pupil dilation with sham stimulation than with the taVNS. Alternatively, the inconsistent findings could reflect the effect of intermixed taVNS and sham in our scapha group, as opposed to the blockwise presentation in Wienke’s study.

Third, we found that across conditions (taVNS and sham) and groups (earlobe and scapha), pupil dilation was largest when the ear was stimulated halfway through the inhalation phase. As we argued above, we believe that this reflects a confound, caused by our finding, previously reported by Schaefer et al. (2025), that baseline pupil size tends to vary across the respiratory cycle. An alternative explanation for the effect of respiratory phase on stimulation-evoked pupil dilation is based on the law of initial values, according to which the size of a physiological response to a stimulus tends to be affected by the prestimulus baseline level of that response system. This law suggests that stimulation-evoked pupil responses might be larger when stimulation is delivered halfway through the inhalation phase, because baseline pupil size is relatively small during inhalation (Figure 3B; Schaefer et al., 2025), offering a greater potential “upward” range. Conversely, stimulation-evoked pupil responses might be smaller during exhalation because the relatively large baseline pupil size during that respiratory phase causes a ceiling effect. Although we cannot exclude this explanation, previous work suggests that the law of initial values does not apply to the pupil system, as long as baseline pupil size is within a natural range (Gilzenrat et al., 2010). In any case, we found no evidence for an enhanced pupil response when taVNS (but not sham) was delivered during the exhalation phase (cf. Sclocco et al., 2019).

Two potential limitations regarding the electrodes should be noted. First, the electrode surface areas were not identical. The clip electrode used for sham stimulation had a surface area of 23.8 mm^2^ per contact. The NEMOS electrode was placed within the curvature of the cymba concha, resulting in an effective contact area of approximately 20–30 mm^2^ per contact, depending on individual differences in the shape of the auricle. In addition, the clip electrode was positioned on the anterior and posterior sides of the auricle, resulting in current passing through the ear tissue. In the case of the upper scapha, this could have led to the unintended activation of intrinsic auricular muscles and other non-vagal afferents, potentially confounding the results (Cakmak, 2019).

## 4. General discussion

In this study, we addressed two key questions. First, how effective is pulsed taVNS at eliciting pupil dilation when active and sham stimulation are delivered intermixed within the same experimental blocks? And second, to what extent does taVNS-elicited pupil dilation depend on the sham location and the respiratory cycle? We draw the following two conclusions.

### 4.1 Feasibility of intermixing taVNS and sham stimulation

Our study is the first taVNS-pupillometry study in which (supra-threshold) pulsed taVNS and sham stimulation were randomly intermixed. When using the earlobe as sham location, we found robust evidence that taVNS increased pupil dilation, in particular when the stimulation duration was 1 second (Experiment 2). This shows that the effect of pulsed taVNS on pupil dilation can be preserved when taVNS and sham are delivered in the same block of trials. The intermixed presentation and 1-second stimulation duration suggest it may be feasible to integrate taVNS into the rapid, event-related task designs of cognitive neuroscientists. However, an important remaining step is limiting the trial duration, which in Experiment 2 ranged from 11 to 13 seconds. When trial durations decrease below 5 seconds, the pupil responses to consecutive taVNS/sham pulses will start to overlap in time. In general, it is possible to estimate the constituent pupil responses using automated model-based deconvolution (Fink et al., 2024; Wierda et al., 2012), or to statistically control for the trial type (and corresponding mean pupil response) on the previous trial. However, the question remains to what extent the effects of taVNS and sham will carry over to the next trial when stimulation pulses are so closely spaced. Future research should address this question.

The effect of taVNS on pupil dilation was much weaker with a stimulation duration of 3.4 seconds (Experiment 1). This is interesting because previous studies with blockwise presentation of active and sham trials but otherwise identical stimulation parameters reported robust taVNS effects on pupil dilation (Lloyd et al., 2023; Sharon et al., 2021), with effect sizes more than twice the size of that obtained in Experiment 1. This discrepancy can possibly be attributed to the intermixing of stimulation types. Although we did not directly contrast the 1-and 3.4-second stimulation durations, we recommend that future studies that want to intermix taVNS and sham stimulation use a relatively short stimulation duration.

### 4.2. Effects of sham location and respiratory cycle call into question anatomical pathway

In Experiment 2, we found strong evidence for an effect of taVNS on pupil dilation in the group in which the sham electrode was attached to the earlobe, the most commonly used sham location. However, we found moderate evidence *against* such an effect in the group that received sham stimulation at the upper scapha. This is potentially problematic for the field, because the discrepancy may relate to criticisms of the use of the earlobe as sham location (Cakmak, 2019): although the earlobe is not innervated by the auricular branch of the vagus nerve, it also differs from the cymba concha in that it has a lower density of sympathetic nerve fibers. Sympathetic fibers are efferent, so they cannot by themselves send signals to the brain. However, the effects of sympathetic activity on local skin conditions, including vasoconstriction and sweat secretion, directly affect how sensory receptors behave (Glatte et al., 2019). This means that adjacent skin areas with different sympathetic density will respond differently to the same stimulus.

It is possible that the separate intensity calibration procedures for the cymba concha and earlobe to some extent control for differences in sensitivity. However, several of our participants spontaneously reported after the experiment that they became more habituated to sham (earlobe) stimulation over the course of the experiment, gradually perceiving it as less intense than the stimulation of the cymba concha. Future research using the earlobe as sham location could benefit from monitoring changes in perceived intensity across the session to ensure consistent stimulation sensations. To avoid this issue, we chose the upper scapha as a sham location. The scapha is not innervated by the auricular branch of the vagus nerve and, as one of the sections of the upper ear, has a similar density of sympathetic nerve fibers as the cymba concha (Cakmak, 2019; Cakmak et al., 2018). Our finding that pulsed stimulation of the cymba concha and scapha elicits a similar pupil response suggests that the pupil response to stimulation of the cymba concha cannot be attributed to activation of the vagus nerve.

Furthermore, if the pupil response to stimulation of the cymba concha would reflect activation of the vagus nerve-NTS-LC pathway, we should have found increased pupil dilation to taVNS but not sham stimulation delivered during the exhalation phase of the respiratory cycle (Sclocco et al., 2019). But our results showed no sign of such a pattern. Instead, taVNS and sham stimulation of the earlobe and scapha all showed a larger pupil dilation response when delivered during the inhalation phase. We argued that this likely reflects a confound produced by a general relationship between pupil size and the respiratory cycle, previously reported by Schaefer et al. (2025) and replicated here. Importantly, this general relationship is unrelated to stimulation of the outer ear. The absence of an exhalation-and taVNS-specific increase in stimulation-evoked pupil dilation does not imply that the NTS and LC are not activated; there are multiple anatomical pathways by which sensory stimulation can activate these brainstem nuclei and elicit a pupil dilation response (Cakmak, 2019; Nieuwenhuis et al., 2011). However, our findings do cast doubt on an important and commonly assumed principle in experimental work using taVNS─namely that the effect of electrical stimulation of the cymba concha on pupil size is mediated by the pathway connecting the vagus nerve to the NTS and LC. Future studies combining taVNS with invasive vagus nerve recordings and pupillometry would be ideally suited to address this question.

In conclusion, the present findings show that pulsed taVNS effects on pupil dilation are preserved when taVNS and sham stimulation of the earlobe are intermixed, at least with a 1-second stimulation duration and a long intertrial interval (11-13 seconds). However, the absence of a taVNS effect when the scapha is used as sham location, as well as the respiration-related findings, question the involvement of the vagus nerve-NTS-LC pathway in pulsed taVNS effects on pupil dilation. These results highlight the need for investigation of pulsed taVNS effects across different conditions and for more direct examination of the underlying neural pathways.

## Acknowledgements

We thank Tristan Nukman and René Tinz for their assistance with data collection.

## Open Practices Statement

This research was not preregistered. The analysis scripts and processed data are openly available on the Open Science Framework (OSF) at https://doi.org/10.17605/OSF.IO/2NJCH. The raw data are available from the corresponding author upon reasonable request.

## Supplementary tables

**Tabel S1.**
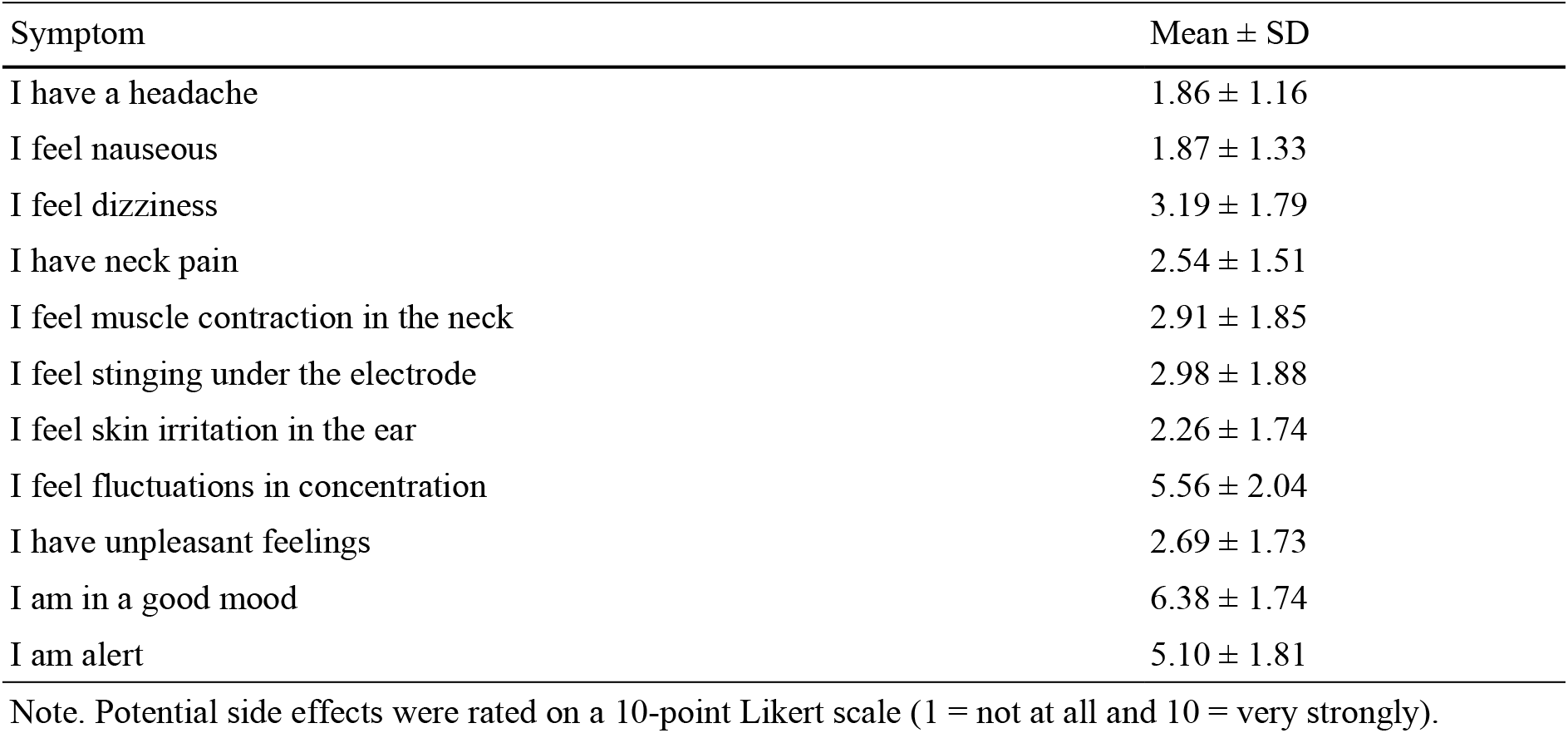
Results of the taVNS aftereffects questionnaire for Experiment 1.

**Table S2.** Results of the taVNS aftereffects questionnaire for Experiment 2.

| Symptom | Earlobe group<br>(M ± SD) | Scapha group<br>(M ± SD) |
| --- | --- | --- |
| I have a headache | 2.38 ± 1.59 | 1.87 ± 1.32 |
| I feel nauseous | 2.17 ± 1.71 | 1.86 ± 1.28 |
| I feel dizziness | 3.27 ± 2.18 | 2.50 ± 1.82 |
| I have neck pain | 2.96 ± 1.98 | 2.25 ± 1.71 |
| I feel muscle contraction in the neck | 3.16 ± 2.19 | 2.26 ± 1.83 |
| I feel stinging under the electrode | 4.03 ± 2.00 | 3.04 ± 2.13 |
| I feel skin irritation in the ear | 2.69 ± 2.20 | 2.19 ± 1.71 |
| I feel fluctuations in concentration | 4.49 ± 2.00 | 5.77 ± 1.58 |
| I have unpleasant feelings | 3.24 ± 2.19 | 2.74 ± 1.77 |
| I am in a good mood | 6.22 ± 2.12 | 5.98 ± 1.98 |
| I am alert | 5.93 ± 2.03 | 5.16 ± 2.07 |

Note. Potential side effects were rated on a 10-point Likert scale (1 = not at all and 10 = very strongly).

